# Discovery and pre-clinical evaluation of antibodies to the NKG2A inhibitory receptor

**DOI:** 10.1101/2022.05.18.492494

**Authors:** Du-San Baek, Ye-Jin Kim, Wei Li, Bernard JC Macatangay, Joshua C. Cyktor, Margaret G. Hines, John W. Mellors, Dimiter S. Dimitrov

## Abstract

NK and CD8^+^ T cells are important cells for cytolysis of cancer cells. The tumor microenvironment can upregulate surface expression on these cells of NKG2A, an inhibitory receptor that can dampen immune responses to cancer leading to immune evasion. To block NKG2A-mediated inhibition, we discovered and characterized two fully human antibodies using phage and yeast display that bind to NKG2A. These antibodies are highly specific for human CD94/NKG2A heterodimer complex, displaying no binding to the activating NKG2C receptor. A mutagenesis study revealed that the serine residue at 170 position (S170) of NKG2A is critical for the selectivity of anti-NKG2A antibodies. In vitro cytotoxic assays showed that NKG2A antibody inhibitors activated primary NK cells and promoted ADCC function of specific antibodies that bind to antigens expressed on cancer cells.

**Summary heading:** Fully human antibodies to the NKG2A inhibitory receptor

## Introduction

The clinical benefit of immune checkpoint inhibitors (ICI) has ushered in a new era of cancer immunotherapy, expanding therapeutic options available to patients with cancer. In particular, monoclonal antibodies (mAbs) targeting programmed-cell death protein 1 (PD-1) and programmed-cell death ligand 1 (PD-L1) have yielded unprecedented, long-lasting benefits to some patients^1-3^. However, because a substantial fraction of patients given anti-PD-1/PD-L1 antibodies do not respond, alternative immuotherapeutic approaches have recently drawn the attention of researchers^4-7^.

One alternative currently being explored is blocking of other inhibitory pathways on NK and T cells that dampen immune responses. NKG2A is an inhibitory receptor of the NKG2 family that has been proposed as a target to complement existing immunotherapies^8-10^. NKG2A forms a heterodimer with CD94 and is dominantly expressed on CD56^+^ NK cells, natural killer T (NKT) cells, and a subset of CD8^+^ T cells. Substantial interaction of CD94/NKG2A heterodimer with its ligand, the non-classical MHC class I molecule HLA-E, leads to the phosphorylation of the immunoreceptor tyrosine-based inhibitory motifs (ITIMs) in the cytosolic domain of NKG2A^11,12^. This event results in transmission of inhibitory signaling and suppression of competing signals from activating receptors, such as NKG2C or D. Elevated expression of HLA-E on the surface of tumor cells or HIV-infected lymphocytes has been reported but NK cells lacking NKG2A eliminate these cells^13-16^. Similarly, inhibition of the NKG2A-mediated signal transmission by an anti-NKG2A monalizumab (humanized Z270 murine mAb) restores NK and T cell effector function in preclinical cancer models^10,13,17^. As a result, monalizumab is now being studied in multiple clinical trials (NCT04307329, NCT04590963, NCT05221840, and NCT05061550) to increase the proportion of patients responding to ICI. The combination of monalizumab with the anti-EGFR antibody, cetuximab, or with the anti-PD-L1 antibody, durvalumab, has led to encouraging responses in a subset of patients with head and neck cancer squamous cell cancer (HNSCC), colorectal cancer (CRC) and non-small cell lung cancer (NSCLC)^13,18-21^. The responses observed to combination mAb therapy are superior to monotherapy with monalizumab (NCT02459301).

In the search for more potent and specific inhibitors of NKG2A, we discovered the first fully human anti-NKG2A mAb designated 1B2 clone. Through engineering with yeast display, we then identified the 1B2-6 clone that exhibits more potent blockade of HLA-E binding to NKG2A. We also show that the 1B2 and 1B2-6 clones activate primary NK cells in PBMCs from healthy donors and promote NK mediated killing of tumor cancer cells by antibodies targeting EGFR and CEACAM5.

## Results

### 1. 1B2 clone selectively binds to NKG2A and inhibits HLA-E interaction

To identify an NKG2A-specific fully human Fab, we produced recombinant NKG2 proteins fused with human IgG1 Fc. Since NKG2A forms a heterodimer with CD94, we expressed four recombinant proteins of NKG2A or 2C with or without CD94 in HEK293F cells to test their use as an antigen (Fig. 1A). We then assessed the impact of CD94 on the binding of monalizumab, a humanized murine Z270 antibody to human NKG2A. We generated a mirroring clone of monalizumab with L234A, L235A, and P329G mutations in the human IgG1 Fc region, abolishing effector function (named mona-IgG in this study) instead of the IgG4 isotype and then tested binding to the four purified NKG2A/C proteins by ELISA. The mona-IgG bound exclusively to CD94/NKG2A heterodimer complex and displayed no binding to the other proteins (Fig. 1B). This result indicated that CD94 is an essential partner domain to generate the binding epitope for monalizumab. Using the validated CD94/NKG2A heterodimer complex, we successfully identified two fully human Fab clones, 1B2 and 2A8, that selectively bound to CD94/NKG2A by ELISA (Fig. 1C). We then investigated whether two Fabs could bind to natural CD94/NKG2A heterodimer on the membrane of primary human NK cells in PBMCs from healthy donors. We confirmed that mona-IgG and 1B2 Fab bound to CD94/NKG2A on primary human NK cells with similar sensitivity of detection (∼20% after subtracting the secondary antibody signal) and that the two antibodies did not bind to NKG2A negative B lymphocyte cell-line (Farage), in flow-cytometry analyses (Fig. 1D, and Fig. S1). By contrast, we did not observe 2A8 Fab binding to CD94/NKG2A on human NK cells in a parallel experiment (Fig. 1D), thus we excluded 2A8 from further investigations. We next sought to assess inhibitory capacity of 1B2 hIgG1 with LALA-PG mutations (named 1B2 IgG) by monitoring the binding of the soluble HLA-E (sHLA-E) ligand to its cognate receptors CD94/NKG2A and CD94/NKG2C. Competitive ELISA results with the tetramer of sHLA-E*01:01(peptide sequence: VMAPRTLVL) demonstrated that mona-IgG and 1B2 IgG inhibited binding of sHLA-E tetramer to CD94/NKG2A in a concentration dependent manner (Fig. 1E). By contrast, two control IgGs did not decrease binding of sHLA-E tetramer to CD94/NKG2C, indicating that the two IgGs (mona-IgG and 1B2 IgG) are NKG2A-specific antibody binders that differentiate between NKG2A and NKG2C, which have only three major amino acid differences (Fig. 1F).

**Figure 1.**
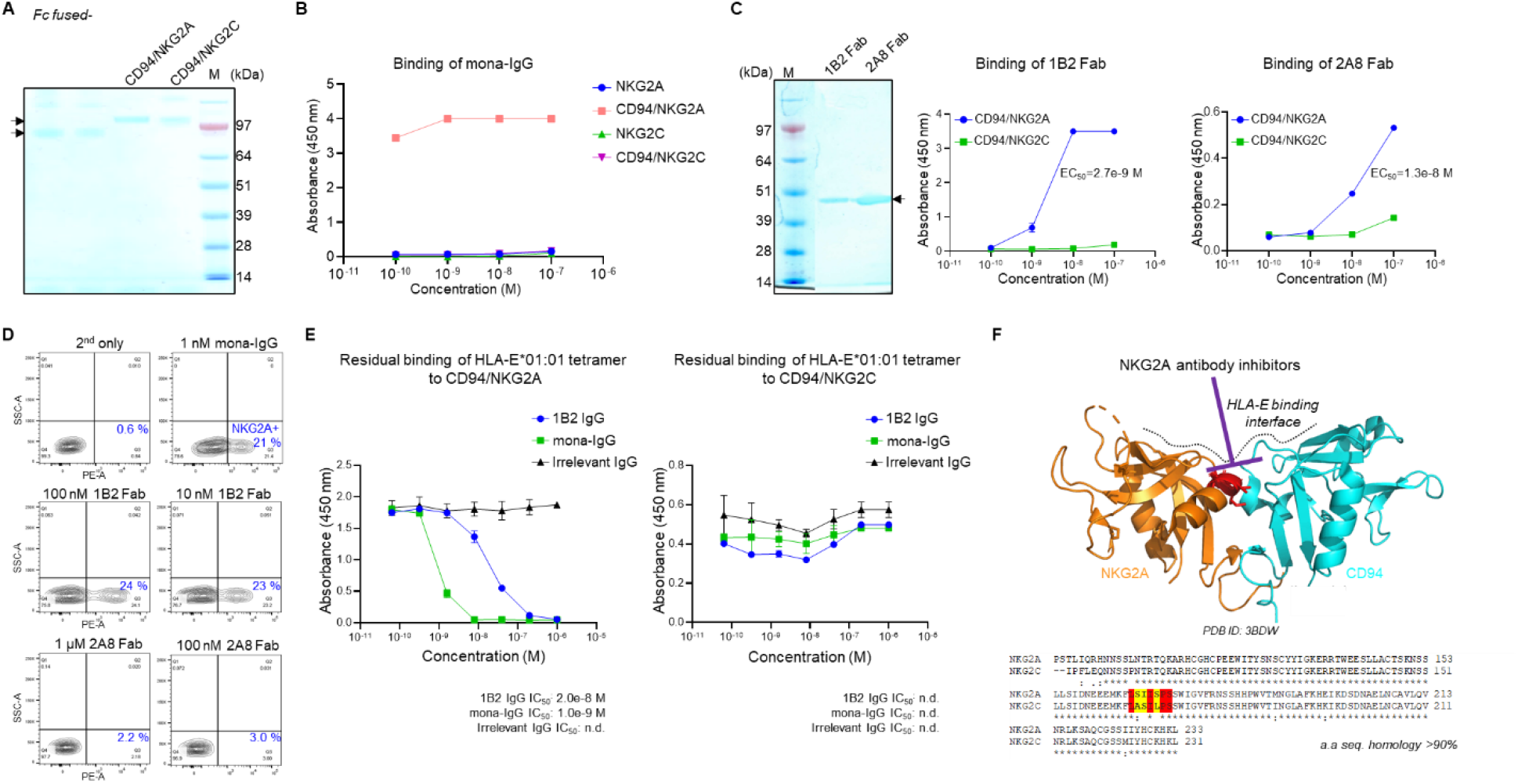
Discovery of NKG2A binding antibody, 1B2 clone. (A) SDS-PAGE gel for purified Fc-fused NKG2A or 2C with or without CD94. Arrows indicate corresponding band for four recombinant proteins (MW: Fc-fused NKG2A or C is 84 kDa, and Fc-fused CD94/NKG2A or C is 109 kDa.) (B) Indirect ELISA result indicating CD94 is essential domain for the binding and specificity of monalizumab analog (mona-IgG). (C) Specificity and binding of purified 1B2 and 2A8 Fab against CD94/NKG2A or 2C in ELISA. (D) Representative density plots depicting percentages of primary NK cells expressing CD94/NKG2A labeled by mona-IgG, 1B2 Fab, and 2A8 Fab in flow-cytometry after treatment of each antibody at indicated concentration for an hour. (E) Competitive ELISA experiment with sHLA-E tetramer demonstrating that 1B2 and mona-IgG selectively block HLA-E binding to CD94/NKG2A. (F) Crystal structure of CD94/NKG2A (PDB ID: 3BDW) and comparison of primary sequences of NKG2A and 2C. Hypothetical critical epitope for 1B2 and mona-IgG is highlighted as red in both structure and sequences and the mismatched residues between NKG2A and C at 167, 168 and 170 positions of NKG2A are colored as yellow. ELISA results are representative of two replicates and data are presented as mean± s.d.

### 2. 1B2 and affinity matured 1B2-6 activate primary human NK cells

Although 1B2 IgG inhibited sHLA-E tetramer binding to NKG2A, the IC_50_ of 1B2 IgG (20 nM) indicated that higher affinity was required for efficient blocking of HLA-E comparable to mona-IgG. Thus, we designed and constructed a single chain Fab (scFab) yeast library for the affinity maturation of 1B2 (Fig. S2). For affinity maturation of VH domain, we applied a strategy to replace 1B2 VH with naïve VHs, with the exception of CDR-H3 and FR4 to avoid major epitope shifting. Similarly, we shuffled 1B2 VL to naïve VLs to generate affinity matured VL. We then monitored binding of scFabs displayed on yeast surface and sorted out mixed yeast library cells that showing improved binding by flow cytometry (Fig. S2). From the library with VH shuffling only, one dominant clone, named 1B2-6, was identified after single clone analysis. This clone exhibited improved blocking of sHLA-E tetramer to NKG2A (Fig. 2C). With the three NKG2A antibodies (mona-, 1B2, and 1B2-6 IgG), we then investigated which residue in NKG2A is involved in selective binding of NKG2A inhibitors. At the amino acid sequence level, three residues at 167, 168, and 170 positions differ greatly among human NKG2 family proteins (A, B, C, E, and H) and in other species (Fig. S3). In addition, these three residues are located at the center position for the interaction with HLA-E in the crystal structure of CD94/NKG2A and HLA-E complex (Fig. 1F), whereas other neighboring residues are almost identical. Thus, we hypothesized that one or two residues may play critical role in epitope generation for NKG2A-specific antibody binders. To test this hypothesis, we constructed three mutant proteins of NKG2A (S167A, I168S, and S170L) and tested the binding of the tree antibodies. Epitope mapping results revealed that a serine residue at 170 position (S170) of NKG2A is critical for the binding of all NKG2A-specific antibodies although the substitution did not lead complete binding loss.

**Figure 2.**
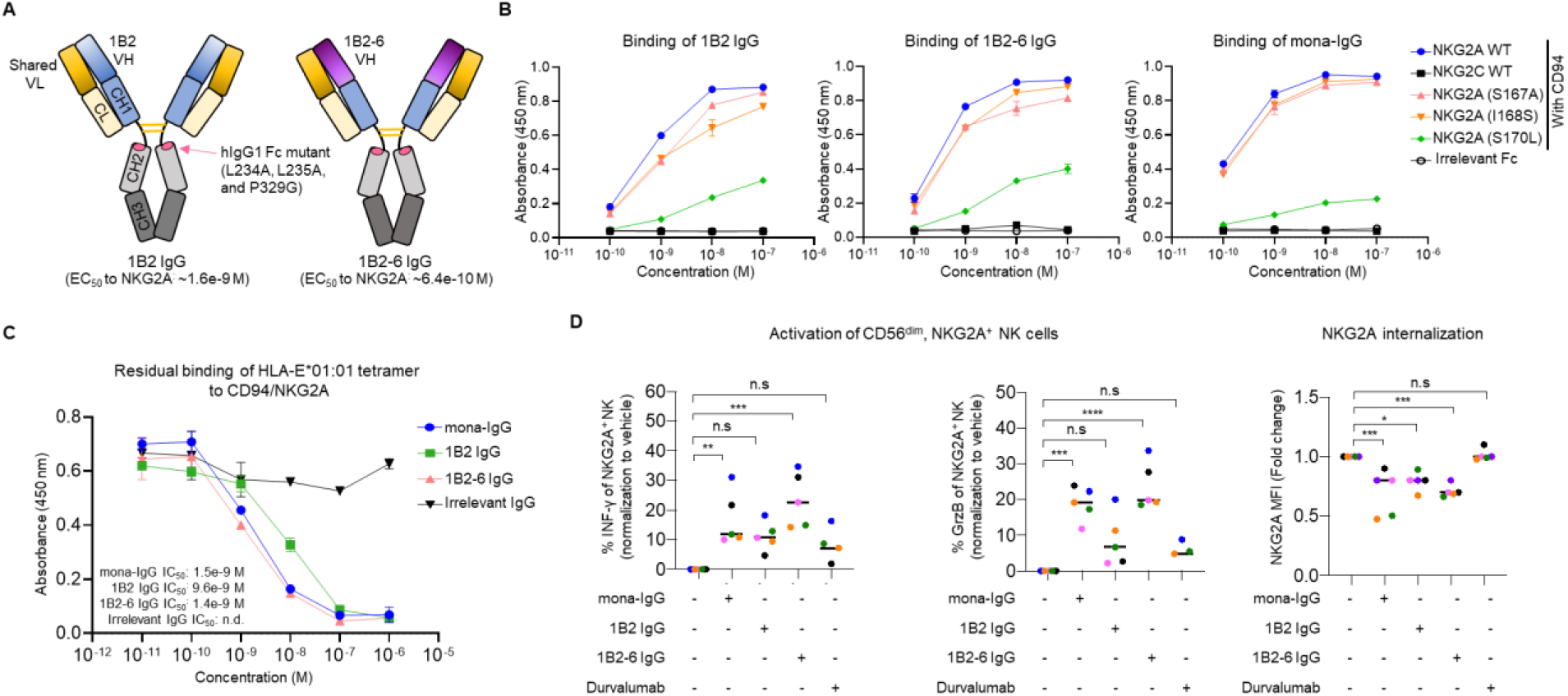
*In vitro* characterization of NKG2A antibody inhibitors. (A) Schematic figure depicting components of 1B2 and affinity matured 1B2-6 human IgG1 having L234A, L235A, and P329G mutations to abolish binding ability to Fc gamma receptors. (B) ELISA experiment for the epitope mapping of three NKG2A inhibitors with mutant proteins of CD94/NKG2A. Each residue at 167, 168, and 170 position of NKG2A was replaced by the indicated NKG2C residue. (C) Competitive ELISA result with sHLA-E tetramer demonstrating affinity matured 1B2-6 IgG has improved blocking capability compared to parent 1B2 IgG. ELISA results are representative of two replicates and data are presented as mean± s.d. (D) Percentage of increased IFN-γ and GrzB level to monitor primary NK cell (CD56^dim^, NKG2A^+^) activation and the decreased surface expression of NKG2A receptor after treatment with NKG2A antibody and anti-PD-L1 antibody, durvalumab, for 24 hours. Each donor is represented by a single dot. Significance was determined by one-way ANOVA, * p < 0.05, ** p < 0.01., *** p < 0.001., n.s means not significant.

We then explored whether NKG2A inhibitors could activate NKG2A^+^ NK cells in either the presence or absence of HLA-E expressing cancer cells. After treatment with NKG2A antibodies for 24 hours, we monitored changes in the cells’ interferon-gamma (IFN-γ) and granzyme B (GrzB) expression level by flow cytometry. The result revealed that NKG2A inhibitor treatment activated CD45^+^, NKG2A^+^ primary NK cells with increased expression of both IFN-γ and GrzB (Fig. 2D and Fig. S4). In addition, NKG2A antibodies led to decreased surface expression of NKG2A on NK cells, although >50% level of initial expression was maintained by re-cycling^22^. The activation of NK cells was not affected by co-culture with HLA-E expressing cancer cells (Fig. S5). Two NKG2A inhibitors, 1B2-6 and mona-IgG, exhibited similar activation levels (>10%). 1B2 IgG also displayed this tendency, but without statistical significance. The order of NK cell activation by NKG2A inhibitors was found to correspond to the order of the IC_50_ values for inhibition of sHLA-E tetramer binding to CD94/NKG2A rather than the degree of NKG2A internalization by NKG2A inhibitors.

### 3. ADCC promoted by NKG2A antibody inhibitors

Previous studies demonstrated that monalizumab promoted antibody-dependent cell-mediated cytotoxicity (ADCC) of the -EGFR hIgG1, cetuximab, with increased CD137 expression^13^. Thus, we investigated whether 1B2 or 1B2-6 IgG could also enhance ADCC by anti-EGFR antibody (cetuximab IgG1) or by anti-CEACAM5 antibody (1G9 IgG1)^23^. Since both antibodies to cancer targets have wild-type Fc region, they bound to Fc gamma receptors while NKG2A inhibitors did not bind due to LALA-PG mutation (Fig. S6). Monotherapy of the three NKG2A antibodies did not lead to NK cell-mediated cytotoxicity (Fig. 3A and 3B). However, combination treatment of NKG2A inhibitors with cetuximab promoted ADCC effects of cetuximab in EGFR^+^/HLA-E^+^ non-small cell lung cancer (NSCLC) A549 and H2030 cells. By contrast, no synergistic effect was observed in EGFR^+^/HLA-E^-^ 293T cells (Fig. 3A). Combination therapy with 1B2 or 1B2-6 IgG promoted the ADCC effect of 1G9 IgG1 compared to 1G9 monotherapy at high doses of CEACAM5^+^/HLA-E^-^ neuroendocrine prostate cancer (NEPC) cell-line, NCI-H660. However, NKG2A inhibitors did not enhance the ADCC effect of 1G9 in CEACAM5-positive prostate cancer Du145-CEACAM5 cells (Fig. 3B). These different effects of combinations were correlated with the different HLA-E expression of each cell line. Interestingly, although 1B2 IgG has been shown to have a lower IC_50_ value in competitive ELISA for inhibition of sHLA-E tetramer binding to CD94/NKG2A, it demonstrated a similar synergistic effect to that shown with 1B2-6 and mona-IgG. These results indicate that the NKG2A-specific fully human antibodies 1B2 and 1B2-6 augment ADCC of tumor-opsonizing antibodies.

**Figure 3.**
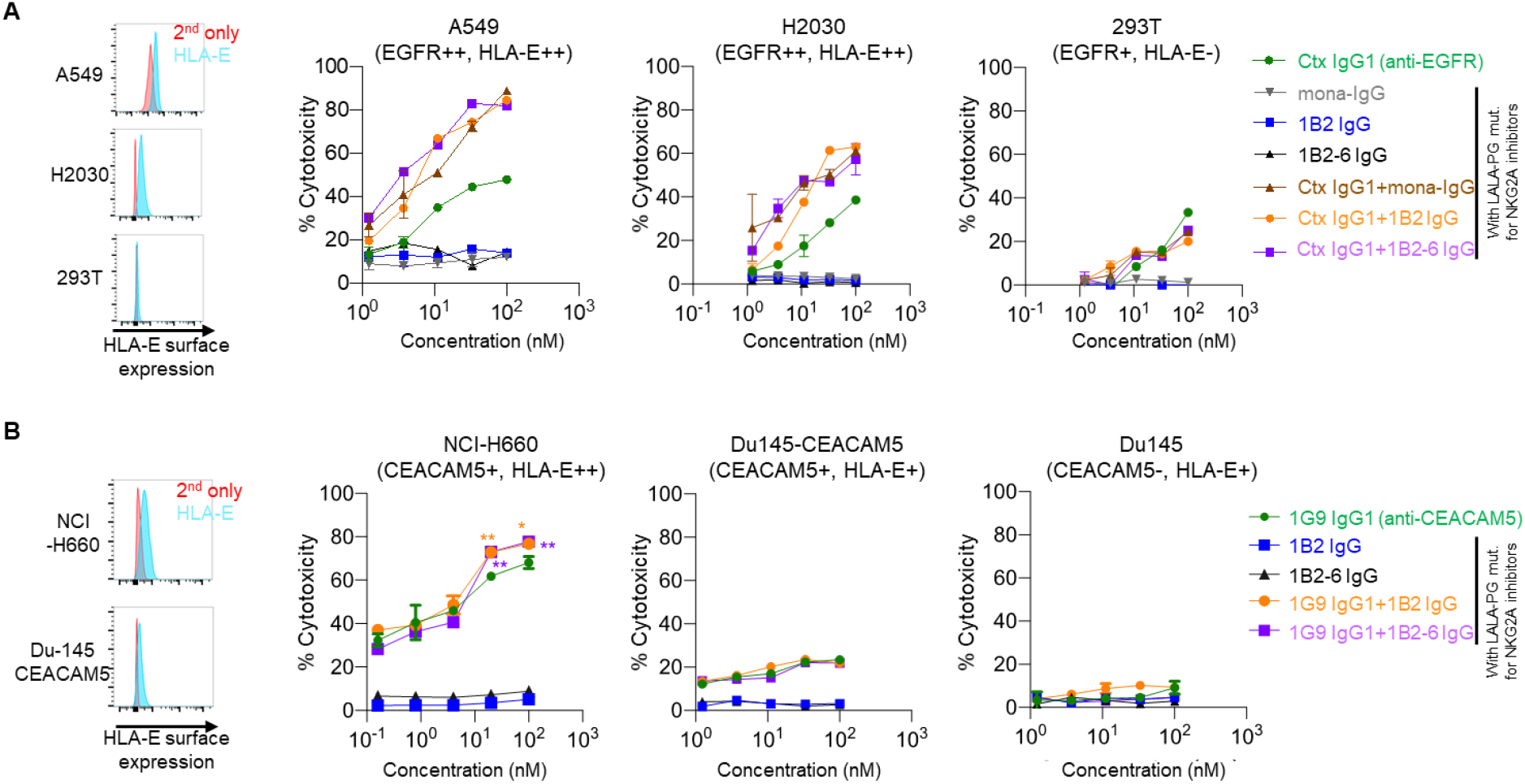
The enhanced ADCC effects of tumor-opsonizing antibodies by combination with NKG2A antibodies *in vitro*. (A) Cell surface HLA-E expression level of A549, H2030, and 293T cells (left panel). ADCC activity of anti-EGFR hIgG1 cetuximab (Ctx) with mona-IgG, 1B2 IgG, or 1B2-6 IgG in NSCLC A549, H2030 cells, and 293T cells (right graphs). (B) Cell surface HLA-E expression level of NCI-H660 and Du145-CEACAM5 cells (left panel). ADCC activity of anti-CEACAM5 hIgG1 1G9 with 1B2 IgG or 1B2-6 IgG in prostate cancer NCI-H660, Du145-CEACAM5, and Du145 cells (right graphs). Right graphs for both (A) and (B) LDH release assay results in presence of primary NK cells from healthy donor PBMCs. E:T ratio, 5:1. Significance was determined by unpaired two-tailed student’s t-test. **, *P* < 0.01; *, *P* < 0.05 vs. Ctx single treatment.

## Discussion

Diverse immune cells continuously eliminate cancerous cells in vivo. This dynamic balance maintains normal tissue function and overall health. Although many immune receptors are involved in the escape or elimination of tumors, the immune checkpoint receptor is now considered a major target to amplify both cytotoxic NK cell and CD8^+^ T cell responses. Following the remarkable success of PD-1/PD-L1 antibody inhibitors, NKG2A inhibitors could become a next generation of immunotherapeutic for cancer. The elevated expression level of CD94/NKG2A often found in tumor infiltrating lymphocytes (TILs) has been previously correlated with decreased survival rate of patients with cancer^13,24^. The subsequently identified interaction between CD94/NKG2A heterodimer with its ligand HLA-E was found inhibit lymphocytes activation. Similarly, aberrant HLA-E display on tumor cells was suggested to cause cancer vaccine resistance; and disruption of NKG2A-mediated inhibitory signaling by NKG2A antibodies potentiated CD8^+^ T cell immunity^25^.

To date, monalizumab is the most widely studied anti-NKG2A antibody due to its evidence of anti-cancer activity in initial clinical trials. However, it is a humanized antibody which still contains murine CDR regions resulting in acute neutralization and clearance by human anti-humanized (murine CDRs) antibodies. In a phase II clinical study in patients with recurrent/metastatic squamous cell carcinoma of the head and neck, anti-monalizumab antibodies were found in 9 patients out of 25 patients (36%)^26^. With competitive panning strategy using NKG2A and NKG2C recombinant proteins, we successfully isolated and characterized first fully human NKG2A inhibitors, 1B2 and 1B2-6 clones, by using both phage and yeast display technologies. The original murine antibody of monalizumab is the Z270 clone, raised from immunized mice injected with CD94^bright^ NK cell clone SA260 (surface phenotype: CD3^−^CD16^+^, CD56^+^, NKp46^+^, NKp44^+^, p70/NKB1^+^, CD94/NKG2A^+^)^17^. Consequently, our investigation showed that monalizumab only bound to CD94/NKG2A heterodimer, not to NKG2A monomer. Our NKG2A inhibitors also did not bind to CD94 lacking NKG2A or 2C monomer (data not shown). Targeting a cooperative conformational epitope formed by CD94 and NKG2A complex allowed anti-NKG2A antibodies to efficiently inhibit HLA-E interaction as HLA-E also binds to a similar interface on CD94/NKG2A^12^. In our mutagenesis study, the serine residue at the 170 position (S170) near the connecting loop between alpha 2 helix and beta 3 sheet of NKG2A domain, was a critical residue for the selectivity of these three NKG2A antibodies. However, substitution of the serine to leucine (S170L) did not lead complete binding loss for all NKG2A inhibitors, indicating that multiple residues cooperatively generate a distinct conformational epitope for these NKG2A inhibitors. Interestingly, at 167, 168, and 170 position (based on human NKG2A numbering), NKG2 family receptors encode mismatched residues even in different species while neighboring residues are identical or similar with the exception of mice. It seems that this region is also critical for the recognition of self HLA-E and shows different binding strength to NKG2 family receptors^12,27^.

In terms of potential clinical relevance, NKG2A antibodies boosted NK cell mediated ADCC in combination with either anti-EGFR or CEACAM5 antibodies. On the other hand, all NKG2A inhibitors failed to elicit NK cell mediated cytotoxicity in the absence of tumor-opsonizing antibody in short-term culture (4 hours). This result is consistent with a previous observations with monalizumab, which also did not improve survival rate of mice when used as a monotherapy^13^. Although disruption of NKG2A inhibitory signaling activates NK cells, the additive anti-tumor efficacy seems to rely on other factors such as epitope and opsonizing properties of the tumor targeting antibody, and involvement of other immune check point receptors, or different glycosylation status of membrane bound proteins^8,28^. As an example, increasing sialylated glycans on tumor cells limited NK cell-mediated immunity including ADCC through the recruitment of sialic acid-binding immunoglobulin-like lectin 7 (Siglec-7)^29,30^. Further investigation is needed to elucidate how NKG2A inhibitors affect CD8^+^ T cell cytotoxic function. In healthy donor PBMCs, the NKG2A^+^ population was below 2% in CD3^+^CD8^+^ T cells. By contrast, the expression level of NKG2A was significantly higher in T cells isolated from cancer patient PBMC samples up to 16% (data not shown). Additional studies may address whether anti-NKG2A antibodies could elicit direct tumor cell killing by T cells and could enhance CDC (define) or ADCP (define) functions by other immune cells. Taken together, the first fully human NKG2A antibodies described here are capable of activating NK cells and augmenting ADCC of tumor opsonizing antibody and thus should be evaluated in preclinical animal models as the next development step.

## Materials and Methods

### Preparation of soluble single-chain CD94 and NKG2A heterodimers

The single chain CD94 (residue 57-179) and either NKG2A (residue 113-233) and NKG2C (residue 111-231) were synthesized in Integrated DNA technologies, Inc. (IDT), and then cloned into pCATDS-Fc plasmid for the mammalian cell expression and purification via *NotI* and *AscI* restriction enzyme sites. The constructed plasmid DNA was amplified from *TOP10F’ E. coli* strain in presence of 100 μg/ml ampicillin and prepared by Midi DNA-prep kit (Macherey-Nagel, 740410) for the transient transfection. Purified DNA was complexed with PEI-Max (Polysciences, 24765-1) and supplied to culture of the Freestyle human embryonic kidney cell-line (Gibco, R79007). 7 days post-transfection, CD94/NKG2A-Fc and CD94/NKG2C-Fc were purified by affinity chromatography with protein A resin (Captiva, NC0997253). Elution of bound proteins to protein A was done by adding 50 mM Glycine buffer pH 3.0 and then buffer was changed to phospho-buffered saline pH 7.4 (PBS) by using PD-10 desalting column (GE, 45-000-148) for the storage. Protein purity was estimated in either SDS-PAGE or size exclusion chromatography (SEC). The concentration of each protein was determined by Nano Drop spectrophotometer 2000C (Thermo, ND2000C).

### ICAT5 Fab library panning against CD94/NKG2A

Fully human Fab phage libraries were constructed previously^23^. ICAT5 and ICAT5-1 phage libraries were pre-blocked with 3% bovine serum albumin (BSA) in PBS (w/v) for 1 h at 25°C. Blocked phages were incubated with 10 nM biotinylated CD94/NKG2A-Fc for 1 h at 25°C in presence of 50 nM CD94/NKG2C-Fc. Bound phages were separated by streptavidin coated magnetic beads (Invitrogen, 11-205-D) and washed 10 times with 1 ml of PBS Ph 7.4 containing 0.1% Tween-20 (w/v). Elution of bound phages was conducted by adding either 1 μM anti-NKG2A antibody or 5 nM HLA-E tetramer. For 2^nd^ and 3^rd^ rounds of panning, reduced concentration of biotinylated CD94/NKG2A (5 nM and 1nM, respectively) and increased competitor ratio of CD94/NKG2C were applied. After 3 rounds of panning, binding of 192 individual clones were analyzed in ELISA and then selected clones were sequenced after plasmid rescue.

### Purification of Fab and IgG (LALA-PG)

The plasmid of positive clones was transformed into *HB2151 E. coli* competent cells, and then colonies were selected in ampicillin containing LB plate (100 μg/ml final concentration) for overnight in incubator at 37°C. The next day, a colony was inoculated in liquid LB + ampicillin media and cultured in 37°C shaking incubator. Isopropyl β-D-1-thiogalactopyranoside (IPTG) was added to the culture at OD_600_ of between 0.4∼0.6 corresponding to around 4×10^8^ cells/ml for a final concentration of 0.1 mM. The culture was relocated to shaking incubator set as 30°C, 200 rpm and incubated overnight. The next day, *E. coli* cells were harvested and resuspended in 1/10 volume of periplasm extraction buffer containing polymyxin B (0.5 mg/ml in PBS pH 7.4) and then incubated on the ice for an hour. Supernatant was collected by centrifugation at 12,000×g for 10 minutes, then loaded into pre-packed Ni-NTA resin. The bound Fab was eluted by adding 300 mM imidazole in PBS pH 7.4 and then imidazole was removed by using PD-10 desalting column. For IgG preparation, IgG cloned plasmid DNA was transfected to HEK239F cells and expressed for 5-7 days post-transfection. Expressed IgG was purified by affinity chromatography with protein A resin. Elution of bound IgG as performed by adding 50 mM glycine buffer pH 3.0 and then storage buffer was changed to PBS pH 7.4 by PD-10 desalting column.

### Variable region shuffled library construction for affinity maturation of 1B2 Fab

The 1B2 single-chain Fab (scFab) was subcloned into the N-terminus of Aga2 in yeast surface-display plasmid pYCAT. Library genes encoding either VL (FR1-CDRL1-FR2-CDRL2-FR3-CDRL3-FR4) or partial VH (FR1-CDRH1-FR2-CDRH2-FR3) with homologous recombination sites were amplified by designed primers (supplementary table X). cDNA isolated from healthy donor PBMCs was used as template DNA. *NheI*/*BsIWI* digested pYCAT plasmid was used for VL shuffled library, and *NheI*/*ApaI* digested plasmid was used for VH shuffling. Transformations were performed by using EZ yeast transformation kit (Zymo research, T2001). With *EBY100* strain sand yeast cells were selected directly in selective SD-CAA media (6.7 g/l yeast nitrogen base, 5 g/l casamino acids, 5.4 g/l Na_2_HPO_4_, 8.56 g/l NaH_2_PO_4_·H_2_O, and 20 g/l dextrose in deionized water) liquid media at 30 °C for O/N. The library screening was performed by three round of fluorescence-activated cell sorting (FACS) using FACS melody (BD Biosciences) against biotinylated CD94/NKG2A-Fc in the presence of a 10-fold molar excess of non-biotinylated CD94/NKG2C-Fc as a competitor. For inducing expression of scFab on yeast surface, 1 × 10^9^ yeast cells were cultured in in SG-CAA media (used galactose instead of dextrose for SD-CAA) at 30 °C for 16h, then induced cells were washed with diluted autoMACS buffer (Miltenyi, 130-091-221) in DPBS pH 7.4. The cell-surface expression and antigen binding level were determined by anti-kappa FITC (Invitrogen, A18854) and streptavidin PE (Invitrogen, S866) or Alexa647 conjugated (Invitrogen, S21374).

### Binding of Fab or IgG in enzyme-linked immunosorbent assay

Binding and specificity of Fab or IgG to CD94/NKG2A or 2C-Fc were analyzed through indirect ELISA. Briefly, Fc-fused proteins were coated on a 96 well plate (Corning, 3690) at 200 ng/well (50 μl volume) in PBS for 2 h at RT. Blocking was carried out with 3% BSA in PBS for overnight at 4°C. The next day, various concentration of Fab or IgG was added to antigen coated plates and incubated for 1 h at 25°C. After three washes, anti-FLAG mouse antibody (M2 clone)-HRP conjugated (Sigma, A8592, 1:3000 dilution) or anti-human kappa goat antibody-HRP conjugated (Invitrogen, A18853, 1:3000 dilution) was added to detect binding of either Fab or IgG. The same volume of 3,3’,5,5’-Tetramethylbenzidine (TMB) (Thermo, PI34028) was added as a substrate and then enzymatic reaction was stopped by adding of 2N sulfonic acid. Biotinylated HLA-E *01:01(peptide sequence: VMAPRTLVL) was purchased from MBL international and it was complexed with streptavidin-HRP in 4:1 molar ratio then 100 pM of tetramer was used for competitive ELISA.

### Cell lines

NCI-H660, Du145, A549, H2030, Farage and 293T cells were purchased from ATCC. Du145 cells were maintained in EMEM (ATCC) supplemented with 10% v/v FBS (Gibco) and 1% penicillin-streptomycin (P/S, Gibco). A549, H2030 and Farage cells were cultured RPMI1640 (ATCC) supplemented with 10% v/v FBS and 1% P/S. 293T cells was maintained DMEM (Gibco) supplemented with 10% v/v FBS, and 1% P/S. NCI-H660 cells were cultured in RPMI 1640 supplemented with 5% FBS, 1X insulin-transferrin-selenium (ITS-G, Gibco), 10 nM hydrocortisone, extra 2 mM l-glutamine, and 1% P/S. Du145-CEACAM5 cells were previously constructed and described^23^. Primary NK cells were cultured with MACS basal NK media with supplementary (Miltenyi Biotec, 130-114-429), 10% v/v human serum (Sigma-Aldrich) and hIL-2 (50 IU/ml, Miltenyi Biotec).

### Flow cytometry analysis

Primary NK cells were enriched from normal human peripheral blood mononuclear cells (PBMCs) (Zenbio Inc.) using NK cell isolation kit (Miltenyi Biotec, 130-092-657). The purity of isolated NK cells was confirmed by staining of APC conjugated anti-human CD56 antibody and PE conjugated anti-human CD16 antibody. The HLA-E expression of target cells was determined by staining of PE conjugated anti-HLA-E antibody. To determine the NKG2A^+^ population of primary NK cells, cells were stained with Alexa488 (AR488)-conjugated anti-NKG2A IgG1. To confirm the cell surface binding of antibodies (mona-, 1B2 IgG, 1B2-6 IgG), cells were treated with antibodies (100 or 10 nM) for 1 h at 4°C and then stained Alexa647-conjugated goat anti-human IgG (Invitrogen, A21445) for 0.5 h at 4°C. For investigation of NKG2A expression level of NK cells, enriched NK cells were activated with 50 IU/ml hIL-2 and treated with the indicated antibodies (100 nM) for 24 h at 37°C. After incubation with antibodies, cells were washed with cold PBS and then the cell surface NKG2A was detected using FITC-or PE-conjugated anti-human CD159a (NKG2A) antibody. For gating of NK cells in the co-culture system with H2030 cells, APC conjugated anti-human CD45 antibody was used. Monoclonal antibodies specific for CD56 (12-0567-42), CD16 (12-0168-42), CD159a (130-113-566), CD45 (17-0459-42), and HLA-E (12-9953-42) were purchased from Thermofisher. For intracellular staining, GolgiPlug containing brefeldin A (Invitrogen, 00-4506-51) was added for the final 4 h of culture to inhibit protein transport from the endoplasmic reticulum to the Golgi apparatus. All intracellular staining was performed using a BD cytofix/cytoperm kit (BD Biosciences, BD554714). Monoclonal antibodies specific for IFN-γ (12-7319-42) and granzyme B (12-8896-42) were purchased from Thermofisher. Durvalumab (anti-PD-L1 antibody, A2013) and cetuximab (anti-EGFR antibody, A2000) was purchased from Selleckchem. Data were acquired using the flow cytometry BD LSR II (San Jose, CA) and analyzed with FlowJo 10.7.1 (Tree Star).

### Functional assay with peripheral human NK cells

The purified human NK cells from PBMCs were used as effector cells. NK cells were incubated with target cell line (H2030) (E:T ratio, 5:1), in presence of 50 IU/ml hIL-2 for 24 h at 37°C. Cells were dispensed into 96-well plates with or without antibodies (monalizumab, 1B2 IgG, 1B2-6 IgG, and durvalumab) at 100 nM for 24 h at 37°C. After treatment of antibodies, cells were fixed and intracellular stained with PE conjugated anti-human IFN-γ antibody and PE conjugated anti-human granzyme B antibody. Cells were washed twice and analyzed on the BD LSR II flow cytometry.

### *In vitro* cytotoxicity assay

The ADCC or cell killing activity of effector cells incubated with anti-NKG2A antibodies with LALA-PG mutation were measured through release of cytosolic LDH from dead target cells using LDH-Glo cytotoxicity assay kit (Promega, J2381). The purified NK cells from normal PBMCs (Zenbio Inc.) were used as effector cells and were incubated with A549, H2030, 293T, NCI–H660, Du145, or Du145-CEACAM5 cells (1×10^4^ cells/well in 96-well plate) as target cells at effector-to-target (E:T) ratio, 5:1 in the presence of the indicated antibodies for 4 h at 37 °C. The percent cytotoxicity was calculated as previously described^23^.

## Acknowledgments and Funding

This work was supported by Pitt internal development funds. We thank other members in Center for Antibody Therapeutics (CAT) for discussion and scientific supports.

## Disclosures

JWM is a consultant to Gilead Sciences and owns shares in Abound Bio and share options in Infectious Disease Connect, unrelated to the current work. DSD owns share in Abound Bio. DSB, YJK, JWM, and DSD are co-inventors of a patent application (US63/289,495).

## Notes

### Competing Interest Statement

The authors have declared no competing interest.

