## Supplemental figures for "Discovery and pre-clinical evaluation of antibodies to the NKG2A inhibitory receptor"

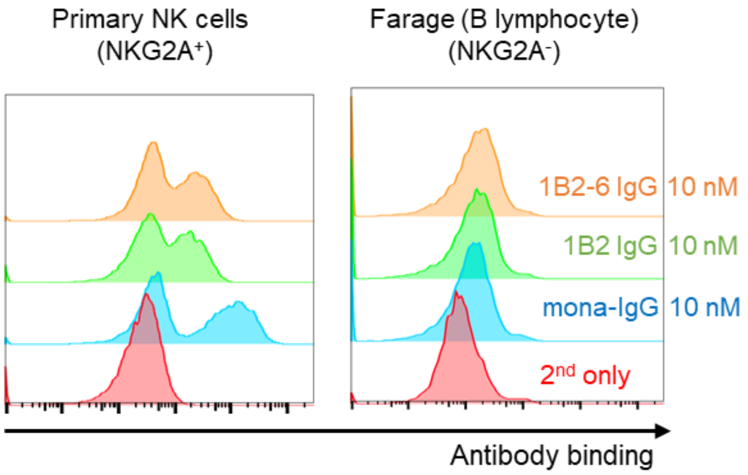


**Figure S1. Binding of antibodies to NKG2A by flow-cytometry.** (A) Binding of mona-IgG, 1B2 IgG, and 1B2-6 IgG antibodies at 10 nM concentration to NKG2A on primary NK cells (A) and the NKG2A negative Farage B cell line (B).

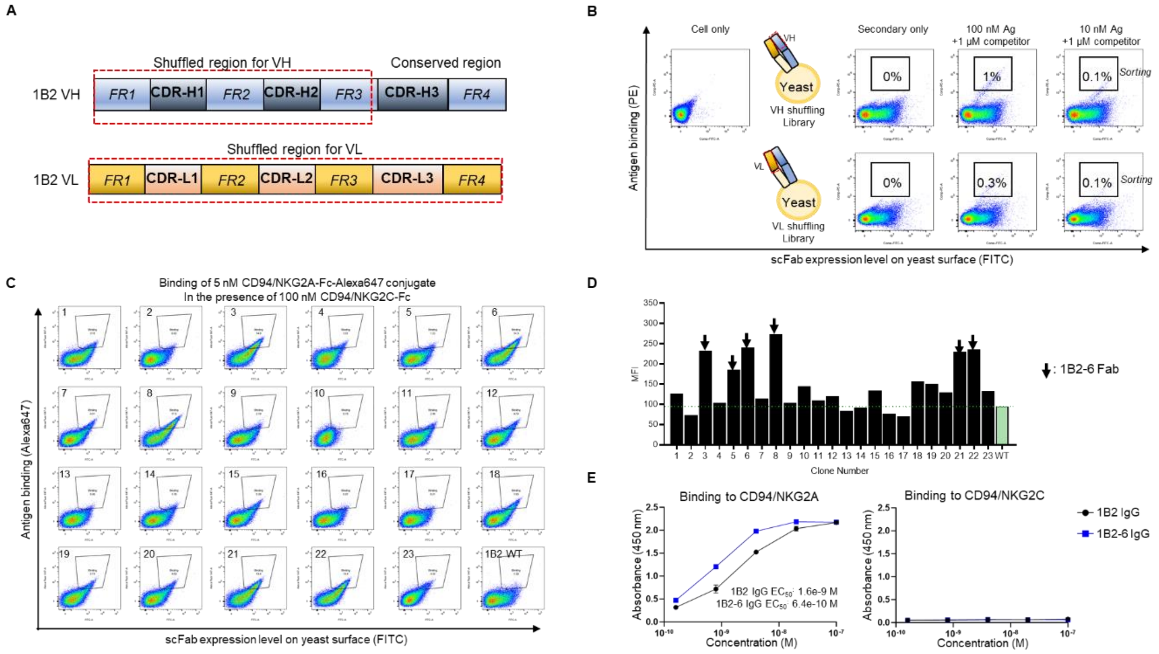


**Figure S2. Affinity maturation of 1B2 Fab by using yeast surface display.** (A) Schematic depicting diversification regions in VH and VL of 1B2. (B) Flow-cytometry analysis of initial VH and VL shuffling libraries to confirm binders are in the constructed scFab library pool and to determine sorting gates. (C) Representative density plots showing scFab expression and antigen binding for individual clones after several rounds of sorting. (D) Bar-graph indicating mean-fluorescence-intensity (MFI) in Panel C. (E) Binding of 1B2 and 1B2-6 IgG to CD94/NKG2A or CD94/NKG2C by ELISA.

A

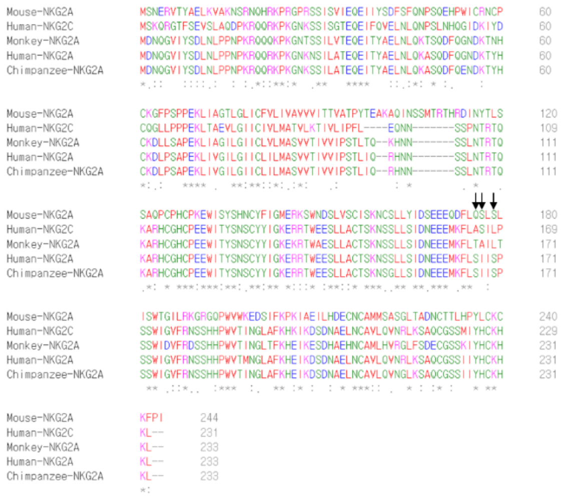

B

| Type of receptor (species) | Name of receptor | Residue at indicated position (arranged to human NKG2A) | | |
|---|---|---|---|---|
| | | 167 | 168 | 170 |
| Inhibitory receptor (Homo sapiens) | NKG2A | Ser (S) | Ile (I) | Ser (S) |
| | NKG2B | Ser (S) | Ile (I) | Ser (S) |
| Activating receptor (Homo sapiens) | NKG2C | Ala (A) | Ser (S) | Leu (L) |
| | NKG2E | Ala (A) | Ser (S) | Leu (L) |
| | NKG2H | Ala (A) | Ser (S) | Leu (L) |
| Inhibitory receptor (Chimpanzee) | NKG2A | Ser (S) | Ile (I) | Ser (S) |
| Inhibitory receptor (Rhesus macaque) | NKG2A | Thr (T) | Ala (A) | Leu (L) |
| Inhibitory receptor () | NKG2A | Gln (Q) | Ser (S) | Ser (S) |

**Figure S3. Primary sequence comparison of the NKG2A inhibitory receptor across different species.** (A) Sequence alignment of NKG2A inhibitory receptors in human, mouse, monkey, and chimpanzee. Human NKG2C sequence is included for comparison. Arrows indicate potentially critical epitopes for NKG2A inhibitory antibody binding. (B) Residues at 167, 168, and 170 position of human NKG2A and corresponding residues in NKG2A family member receptors in different species. Green color indicates hydrophilic residues and red color highlights hydrophobic residues.

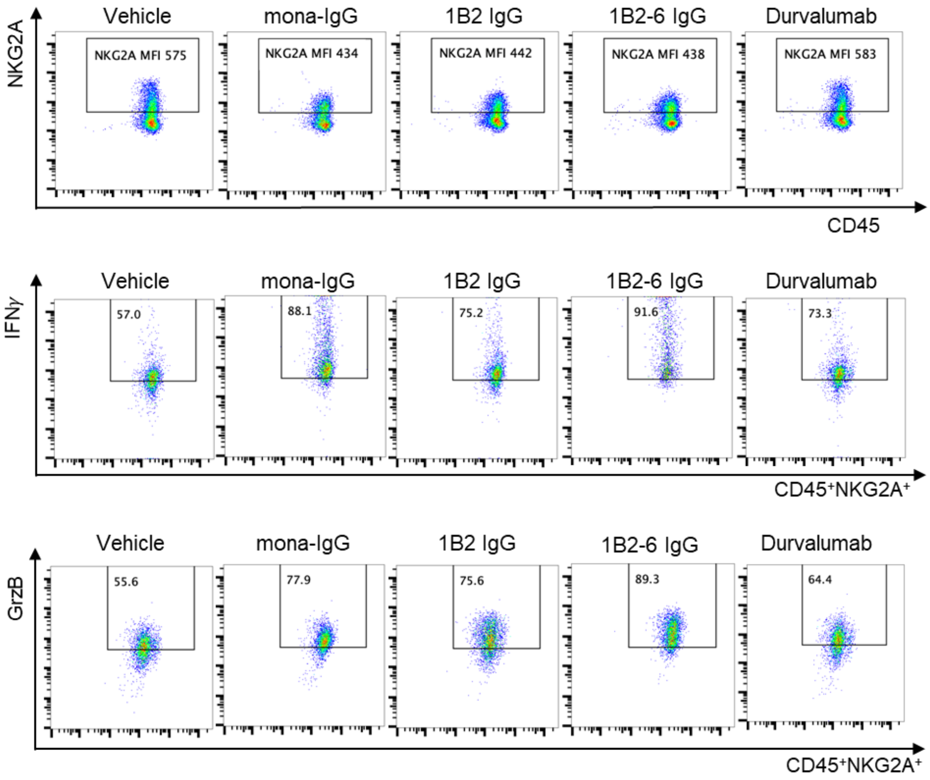


**Figure S4. Assessment of NKG2A surface expression and IFN-γ, GrzB levels by flow-cytometry.** Representative density plots indicating percentage of cells expressing IFN-γ and GrzB to assess NK cell (CD56$^{dim}$, NKG2A$^{+}$) activation, and decreases in surface expression of NKG2A receptor after treatment with anti-NKG2A antibody compared with anti-PD-L1 antibody (durvalumab) for 24 hours.

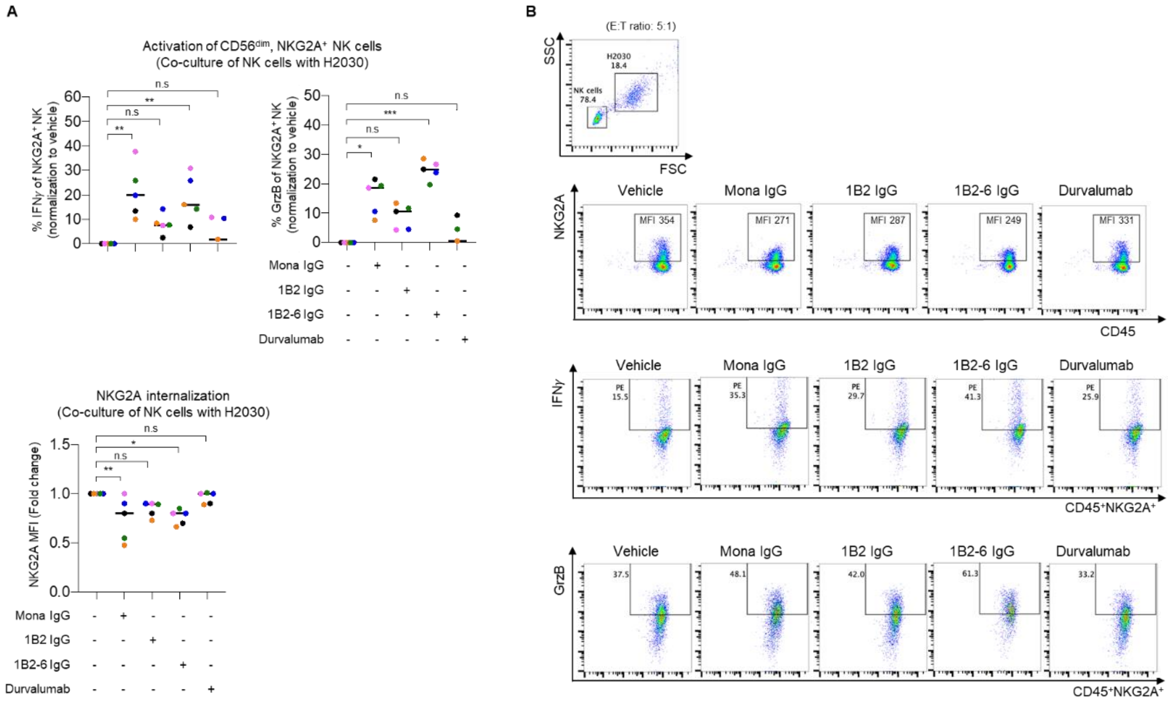


**Figure S5. Assessment of NKG2A surface expression and IFN-γ, GrzB levels by flow-cytometry during co-culture with HLA-E$^{+}$ cancer cells H2030.** (A) Percentage of cells expressing IFN-γ and GrzB to assess primary NK cell (CD56$^{dim}$, NKG2A$^{+}$) activation and decreases in surface expression of NKG2A receptor after treatment for 24 hours with anti-NKG2A antibodies and the anti-PD-L1 antibody, durvalumab. Each donor is represented by a single dot. Significance was determined by one-way ANOVA, * $p < 0.05$, ** $p < 0.01$., *** $p < 0.001$., n.s means not significant. (B) Representative density plots and gate scheme for Panel A.

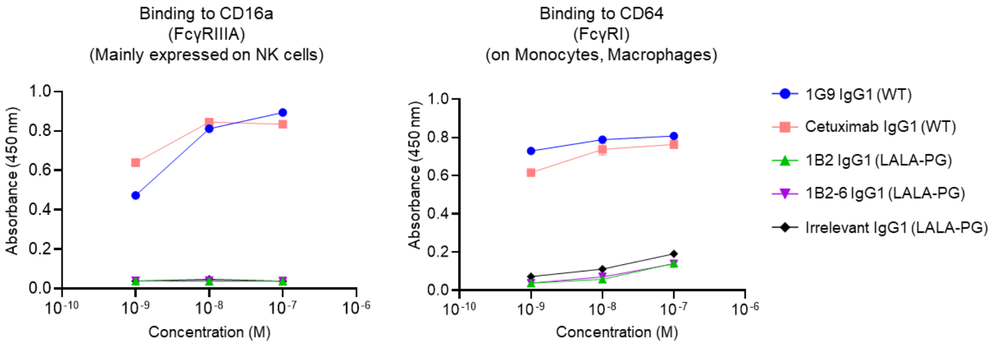


**Figure S6. Binding of anti-NKG2A antibody to recombinant CD16a and CD64.** With L234A, L235A, and P329G mutations in human IgG1 Fc region (LALA-PG), anti-NKG2A antibodies show decreased binding to both CD16a and CD64 in ELISA, compared to IgG1 wild-type (WT).
